# Bursting firing in ventromedial hypothalamic neurons exerts dual control of anxiety-like behavior and energy expenditure

**DOI:** 10.1101/2020.12.20.423695

**Authors:** Jie Shao, Dashuang Gao, Yunhui Liu, Shanping Chen, Nian Liu, Lu Zhang, Xinyi Zhou, Qian Xiao, Liping Wang, Hailan Hu, Fan Yang

## Abstract

Exposure to chronic stress induces anxiety-like behavior and metabolic changes in animals, resulting in adaptive or maladaptive responses to the stressful environment. Recent studies have indicated the dorsomedial ventromedial hypothalamus (dmVMH) as an important hub that regulates both anxiety and energy homeostasis. However, up to now, how dmVMH neurons exert dual control of chronic stress-induced anxiety and energy expenditure remains poorly understood. Here, we established a chronic-stress mouse model that induced anxiety-like behavior, reduced food consumption, and decreased energy expenditure. We found that *c-fos* expression increased and theta band power is higher in the dmVMH after chronic stress, and the proportion of burst firing neurons significantly increased, which was mediated by elevated expression of T-type calcium channel Cav 3.1. Optogenetically evoked burst firing of dmVMH neurons induced anxiety-like behavior, shifted the respiratory exchange ratio toward fat oxidation, and decreased food intake, while knockdown of Cav3.1 in the dmVMH had the opposite effects. Collectively, our study first revealed an important role of dmVMH burst firing in the dual regulation of anxiety-like behavior and energy expenditure, and identified Cav 3.1 as a crucial regulator of the activity of the burst firing neurons in dmVMH. These molecular and cellular level findings will advance our understanding of the chronic stress-induced emotional malfunction and energy expenditure disorders.

## Introduction

Exposure to stressors regulate neural activity that integrate the aversive behavior and energy expenditure to facilitate stress coping and survival^1–3^, however long-term stress causes adverse effects, impairing both mental and physiological functions ^4,5^ including anxiety-like behavior ^6,7^ and metabolic malfunctions in glucose ^8,9^ and lipid tissues ^3,10^. Particular effort has been expended to identify the crucial central nodes that interconnect the regulation of emotion and metabolic processes ^3,9,11^. The hypothalamus, especially the ventromedial hypothalamus (VMH), has been found to play crucial roles in controlling both innate survival behaviors ^12,13^ and energy homeostasis ^11,14,15^.

The VMH is an evolutionarily conserved deep subcortical nucleus^16^. While the ventrolateral part (vl) of the VMH modulates a series of social behaviors ^17–19^, the dorsomedial part (dm) is specifically involved in maintaining energy homeostasis ^14,15,20,21^ and stress response^13,22^. Oscillations in the dmVMH can act on sympathetic excitation and regulate energy expenditure ^23,24^. Steroidogenic factor-1 (SF-1)-expressing neurons are enriched in the dmVMH and play important roles in controlling food intake and energy expenditure ^14,15,20,21^. Optogenetic activation of VMH SF-1 neurons induces aversive behavior in mice ^13,22^. Furthermore, SF-1 neurons can act as a nutrient-sensitive switch between feeding and anxiety states when facing potential stresses ^2^. Given the complicated functions of SF-1 neurons ^19^, it is important to dissect how molecular or electrophysiological distinct neuronal subtypes regulate emotional state and energy homeostasis. However up to now, how dmVMH neurons control neuronal firing to regulate chronic stress-induced anxiety and energy metabolic disorders is elusive.

Recent studies have demonstrated the role of burst firing neurons in regulating emotional state and brain functions; neuronal burst firing and oscillation are critical for the transmission of information and regulation of specific physiological processes in crucial brain areas^25–30^. In the hippocampus, burst firing of CA1 pyramidal neurons play an important role in regulating N-methyl-d-aspartate (NMDA)-mediated transmission and the epileptogenic mechanism ^25^. In the ventral subiculum, chronic social defeat stress increases both T-type calcium channel (T-VGCC)-mediated currents and expression of Cav3.1 protein ^29^. In the lateral habenula, NMDA-receptor-and-T-VGCC-dependent bursting activity are increased after chronic stress and, most strikingly, can drive depression-like behaviors ^26^. Based on these evidences, we decided to investigate the specific function of bursting firing in the dmVMH, and whether it may be involved in chronic stress-induced anxiety-like behavior and energy metabolic changes.

In this study, we first identified that dmVMH neurons can be classified into silent, tonic-firing, and bursting subtypes. We also established a chronic-stress mouse model that induced anxiety-like behavior, reduced food consumption, and decreased energy expenditure. We found a significantly increased proportion of bursting neurons under chronic stress conditions, which was mainly caused by elevated T-type calcium channel Cav3.1 expression. Importantly, optogenetically-induced dmVMH burst firing drive anxiety-like behaviors, shifted respiratory exchange ratio and decreased average energy expenditure in naïve mice. Conversely, knockdown of Cav3.1 in the dmVMH rescued chronic stress-induced anxiety-like behavior and energy expenditure changes. Taken together, our study first identified Cav3.1-mediated burst firing pattern in the dmVMH, and demonstrated its crucial role in regulating anxiety-like behavior and energy expenditure during the chronic stress.

## Results

### Chronic stressors induced anxiety-like behavior and energy expenditure change

Repeated exposure to stressors modifies the activity of neuronal circuits associated with a variety of abnormalities, including behavior, emotional state, and energy expenditure ^2^, which adaptively or maladaptively respond to the stressful situation. To probe the impact of chronic stress on behavior and energy metabolism, we applied unpredictable chronic stressors to establish a stress mouse model. During the chronic stress period, mice were exposed to random stressors for four weeks, as illustrated in Fig. 1a, with no stressors used in the control group. Both the wild-type and stress groups were tested in a comprehensive phenotyping paradigm. Consistent with previous reports ^6,7^, we found that chronic stress induced a state of anxiety (Fig. 1b-e). Compared with the control group, mice in the stress group spent less time in the central area of the open field and open arm of the elevated plus maze without any change in locomotion distance (Fig. 1c and e). The number of entries into the central area and open arms also decreased in the stress group compared with the control group (Fig. 1c and e).

**Fig 1.**
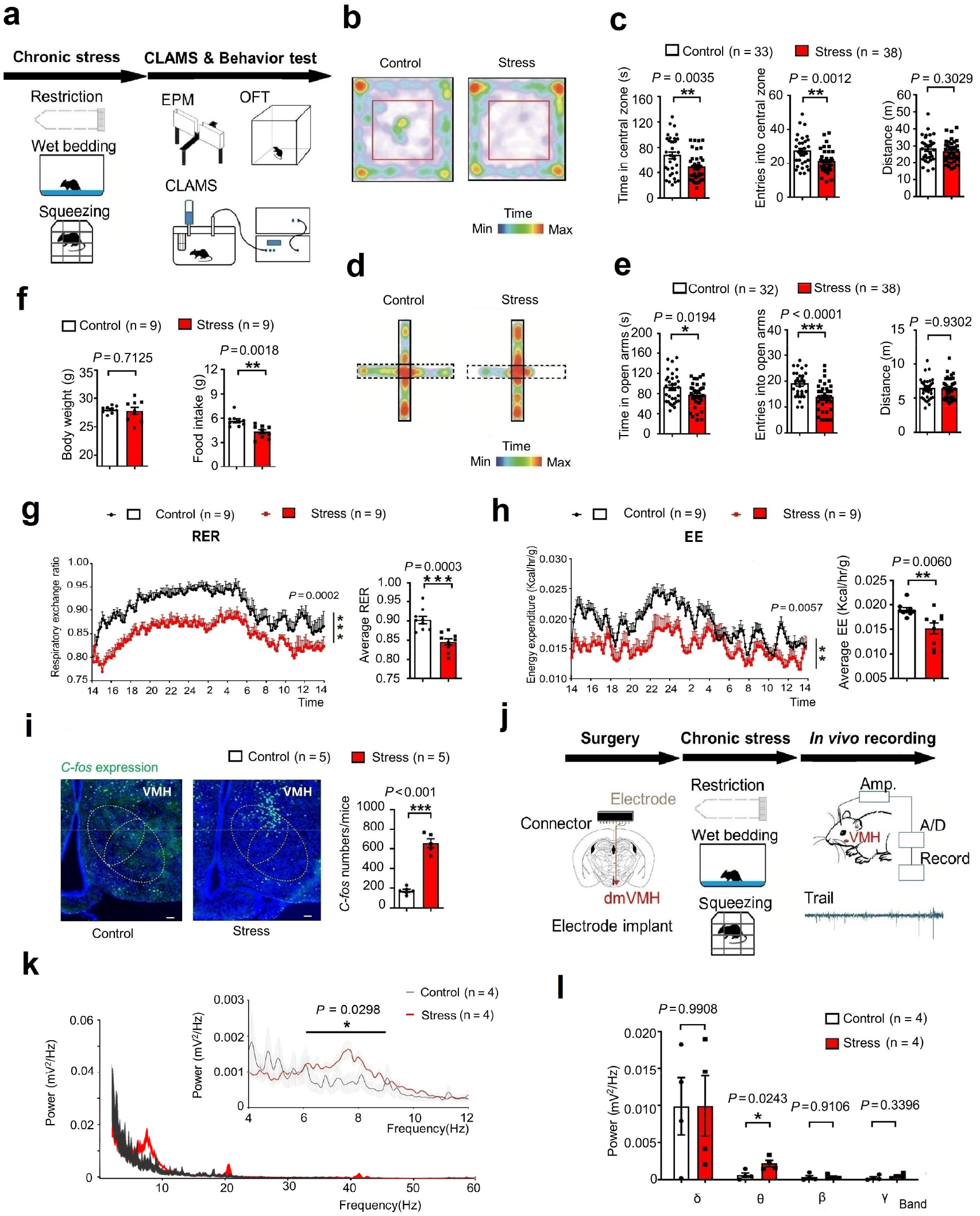
Chronically stressed mice exhibited anxiety-like behavior, altered metabolism, and *in vivo* electrophysiological dmVMH activity. **(a)** Illustration of unpredictable chronic stress protocol and phenotype assessment, with one stressor treatment randomly chosen per day, lasting for four weeks. **(b)** Residence time in each site of open field (blue, less time; red, more time). **(c)** Behavioral analysis of control (n = 33) and stress group mice (n = 38) in open field test showed significant decrease in both time spent in central area (unpaired Student’s *t*-test, *P* = 0.0035) and entries into central area (unpaired Student’s *t*-test, *P* = 0.0012), but no obvious change in travelling distance (unpaired Student’s *t*-test, *P* = 0.3029). **(d)** Residence time in each site of elevated plus-maze (blue, less time; red, more time) of control and stress groups. **(e)** Control (n = 32) and stress groups (n = 38) in elevated plus-maze showed significant decrease in time spent in open arm (unpaired Student’s *t*-test, *P* = 0.0194) and entries into open arm (unpaired Student’s *t*-test, *P* < 0.0001), but no obvious change in locomotion (unpaired Student’s *t*-test, *P* = 0.9302). **(f)** Body weight monitored after chronic stress period showed no obvious change compared with naïve mice (n = 9 in each group, unpaired Student’s *t*-test, *P* = 0.7125), and 24-h food intake after overnight fasting decreased in stress group (n = 9 in each group, unpaired Student’s *t*-test, *P* = 0.0018) compared with control group (n = 9). **(g)** Average respiration exchange ratio (RER) decreased in stress group compared with control group (unpaired student’s *t*-test, *P* = 0.0003, n = 9 mice in each group); RER curve shifted after chronic stress (two-way ANOVA, *P* = 0.0002, *F*(1, 16) = 21.95). **(h)** Significant decreases in 24-h energy expenditure (EE) curve and average EE were observed in stress group (two-way ANOVA, *P* = 0.0057, *F*(1, 16) = 20.2; unpaired Student’s *t*-test, *P* = 0.0060). **(i)** Increased *c-fos* expression in dmVMH under chronic stress (unpaired Student’s *t*-test, *P* < 0.001, n = 5 mice in each group). **(j)** Schematic of electrode implantation and *in vivo* electrophysiology recordings. **(k** and **l)** Power spectral density of local field potential (LFP) in dmVMH after chronic stress, with significant power improvement observed in theta band (two-way ANOVA, *P* = 0.0298, *F*(1, 14) = 5.845,; unpaired Student’s *t*-test, *P* = 0.0243, n = 4). Data are means ± SEM. * *P* < 0.05, ** *P* < 0.01, *** *P* < 0.001. CLAMS, comprehensive laboratory animal monitoring system.

We next explored whether chronic stress affected energy expenditure in stressed mice. The body weight of each mouse was measured after the chronic stress period, and no significant difference was observed between the control and stress groups (Fig. 1f). Both the respiratory exchange ratio (RER) and food intake were monitored for 24 h using comprehensive laboratory animal monitoring system (CLAMS) cages. Results showed that the average RER of the stress group was much lower than that of the control group, which represented a shift toward fat oxidation (Fig. 1g). Energy expenditure was calculated from oxygen consumption and RER. Average energy expenditure also showed a significant decrease in stressed mice (Fig. 1h). To determine the underlying cause of different RER in the treatment groups, we also analyzed food intake. As expected, caloric intake in stressed mice was significantly lower than that in the controls, consistent with the known effects of anxiety on food intake (Fig. 1f). Taken together, these findings suggest that chronic stress can induce anxiety-like behavior, reduce food consumption, and shift the RER toward fat oxidation.

Given its important role in regulating anxiety, feeding, and energy expenditure ^2,21^, we hypothesized that the dmVMH may be involved in the above chronic stress-induced changes. After chronic stress, mice were perfused, and *c-fos* expression was determined via immunostaining. Higher *c-fos* expression was found in the dmVMH of chronically stressed mice than in that of the control group (Fig. 1i), suggesting that more dmVMH neurons were activated under chronic stress conditions. We then employed *in vivo* electrophysiology to record neuronal activity in the dmVMH. Mice were implanted with Ni-Cr electrodes into the dmVMH after chronic stress induction (Fig. 1j), and dmVMH neuronal activity was monitored after recovery from surgery. The local field potential (LFP) results demonstrated that stressed mice exhibited higher theta band power in the dmVMH than that of control mice (Fig. 1k-l). Together with the *c-fos* staining results, our data suggest that dmVMH neuronal activity is significantly changed after chronic stress.

### Chronic stressors induced burst firing in dmVMH

The above experiments revealed enhanced neuronal activity in the dmVMH under chronic stress, but the underlying mechanism was unclear. To explore which electrophysiological characteristics of dmVMH neurons changed after chronic stress, we recorded 84 dmVMH neurons from 38 stressed mice and 85 dmVMH neurons from 32 control mice under whole-cell patch clamp configuration and analyzed the electrophysiological characteristics. Our data demonstrated that these neurons displayed shorter onset time and depolarized RMP, on average, compared with the control group (Fig. 1a). The alterations implied that the overall excitability of dmVMH neurons was enhanced under chronic stress, consistent with the *c-fos* immunostaining and *in vivo* electrode recording results.

We carried out a comprehensive analysis of the membrane properties of dmVMH neurons to quantitatively determine their electrophysiological diversity (Fig. 2b). To classify these neurons, we measured and analyzed eight electrophysiological parameters (Table 1) and performed cluster analysis. The resulting dendrogram in Fig. 2b (with rescaled distance shown along the vertical axis) illustrates the similarity between clusters. Furthermore, cluster analysis showed that all neurons could be divided into three subtypes: i.e., silent, tonic-firing, bursting. These three dmVMH neuronal subtypes displayed distinct electrophysiological properties (Fig. 2b, Extended Data Fig. 1).

**Table 1.** Electrophysiological properties of three dmVMH neuronal subtypes.

|  | Silent<br>(n = 19) | Tonic-firing<br>(n = 38) | Bursting<br>(n = 28) |
| --- | --- | --- | --- |
| Input resistance (MΩ) | 535.30 ± 21.83 | 661.72 ± 19.84 | 662.72 ± 24.21 |
| Onset Time (ms, 50 pA) | 37.17 ± 2.53 | 18.20 ± 1.28 | 25.84 ± 1.87 |
| RMP (mV) | -65.27 ± 1.08 | -57.50 ± 0.92 | -63.39 ± 1.07 |
| Overshoot by H-current<br>(mv, -100 pA) | 7.26 ± 1.17 | 10.71 ± 0.94 | 12.01 ± 1.19 |
| Evoke firing rate (Hz, 100<br>pA) | 17.53 ± 0.60 | 21.76 ± 0.50 | 22.00 ± 0.73 |
| AHP (mV) | -12.38 ± 0.53 | -15.2 ± 0.50 | -14.26 ± 0.65 |
| Half width (ms) | 2.17 ± 0.08 | 2.00 ± 0.04 | 2.04 ± 0.05 |
| AP amplitude (mV) | 71.79 ± 1.88 | 73.29 ± 1.25 | 76.27 ± 1.28 |

Data are represented as mean ± SEM. RMP: resting membrane potential; H-current: hyperpolarization-activated current; AHP: after-hyperpolarization potential; AP: action potential.

**Fig. 2.**
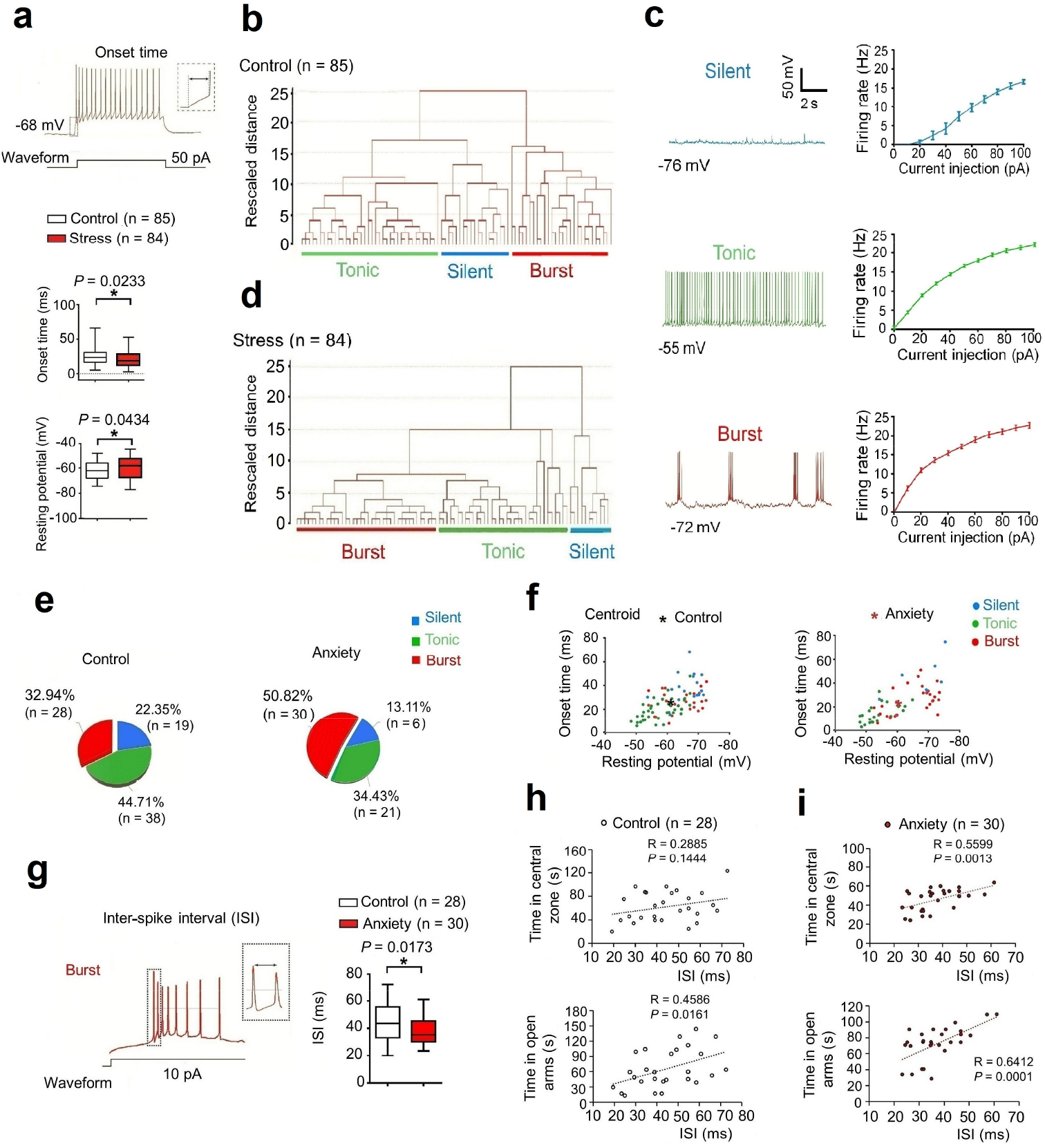
Chronic stress-induced enhancement of burst firing in dmVMH neurons. **(a)** Increased average onset time (unpaired Student’s *t*-test, *P* = 0.0233) and depolarized average resting membrane potential (RMP) (unpaired Student’s *t*-test, *P* = 0.0434) in 84 dmVMH neurons from stressed mice compared with 85 neurons from wild-type mice. The box plotted at the median extending from the 25-75th percentile, and the whisker represents Min to Max distribution. **(b)** Cluster analysis of 85 dmVMH neurons from 35 normal mice. Dendrogram of cluster analysis shows that dmVMH neurons could be classified into three subtypes: i.e., silent, tonic-firing, and bursting. **(c)** Electrophysiological properties of silent, tonic-firing, and bursting dmVMH neuronal subtypes. left: whole-cell recording traces of three neuronal subtypes without current injection; right: frequency-current curve of three subtypes at current injections of 0 to 100 pA and 10 pA/step. **(d)** Cluster analysis of 84 dmVMH neurons from 39 stressed mice. Dendrogram of cluster analysis shows these dmVMH neurons can be classified into three subtypes: i.e., silent (n = 13), tonic-firing (n = 35), and bursting (n = 38). See also Figure S1 and S2. **(e)** Pie chart of percentages of neuronal dmVMH subtypes in control group and stressed mice with obvious anxiety-like behavior (anxiety group). **(f)** Distribution of 85 dmVMH neurons in control mice (left) and 57 dmVMH neurons in anxiety group (right) using onset time-RMP coordinate system, with coordinate of centroid (★) indicating average onset time and RMP. Blue, silent; Green, tonic-firing; Red, bursting. Centroid coordinate was determined by average onset time and RMP. Shift in centroid after chronic stress represents shorter average onset time and more depolarized average RMP, caused by changes in proportion of three subtypes. **(g)** Inter-spike interval (ISI) of bursts in dmVMH neurons of control and stressed mice. Left, Example of burst firing and ISI; right, ISI of burst firing dmVMH neurons (n = 30) in anxiety group decreased significantly compared with that in control group (n = 28, unpaired Student’s *t*-test, *P* = 0.0099). **(h)** ISI of dmVMH bursting neurons in control group is slightly correlated with the residing time in open arms of EPM, but not with the time spent in central area of open field (n = 28 cells from 21 mice). **(i)** ISI of dmVMH bursting neurons in stressed mice which displayed obvious anxiety behavior is significantly correlated with both the residing time in open arms and the time in central area of open field (n = 30 cells from 20 mice). Data are means ± SEM * *P* < 0.05, ** *P* < 0.01.

We then applied cluster analysis using the same methods as depicted above (Fig. 2d) to explore overall changes in the electrophysiological properties of dmVMH neurons after chronic stress. We found that dmVMH neurons in stressed mice could also be divided into three subtypes. Onset time, RMP and recording site of these three subtypes were also compared with those of the control group (Extended Data Fig. 2). By analyzing data from behavior tests and electrophysiological recordings, we found the proportion of burst firing neurons in stressed group, especially in mice with obvious anxiety-like behavior (anxiety group), increased significantly compared with that in the control group, whereas the proportion of the other subtypes decreased (Fig. 2d-e). This alteration could explain why dmVMH neurons from stressed mice demonstrated a shorter average onset time of action potentials induced by 50 pA current injection (Fig. 2a), as burst firing neurons displayed a similar onset time as tonic-firing neurons but a shorter onset time compared with silent neurons (Fig. 2f, Extended Data Fig. 2b). In addition, the increased proportion of bursting neurons and more depolarized tonic-firing neurons (Fig. 2f, Extended Data Fig. 2a) may have contributed to the higher average RMP in dmVMH neurons after chronic stress (Fig. 2a). Furthermore, the average inter-spike interval (ISI, between first and second spike of a burst) of dmVMH burst firing neurons in anxiety group was shorter than that of the control group (unpaired Student’s *t*-test, *P* = 0.0173), and a shorter ISI represents higher frequency firing in a single burst (Fig. 2g). We also found that ISI of dmVMH bursting neurons was more correlated with the time in central area or open arms in mice of stress-induced anxiety compared with the mice in control group (Fig. 2h,i). Taken together, these results demonstrate a possible link between increased bursting activity in the dmVMH and chronic stress induced anxiety.

### Optogenetic manipulation of burst firing neurons in dmVMH

Given the enhancement of burst firing in the dmVMH after chronic stress, we investigated whether induced burst firing of dmVMH neurons alone in unstressed mice was sufficient to produce a similar response in behavior and energy metabolism. We first applied optogenetic manipulation to elicit a low-threshold spike and mimic the generation of bursting in dmVMH neurons (Fig. 3a). *In vitro* patch clamp experiments on brain slices indicated that yellow light illumination at 0.1 Hz could slightly hyperpolarize and depolarize bursting neurons periodically, which enabled the re-activation of calcium channels and generation of burst firing during the light “ON-OFF” intervals in bursting neurons (Fig. 3b). The evoked burst firing did not follow the “ON-OFF” intervals when the illumination frequency was over 0.1 Hz (Fig. 3b). In addition, illumination at 0.1 Hz exerted no significant effects on the tonic-firing or silent neuronal subtypes (Fig. 3c).

**Fig. 3.**
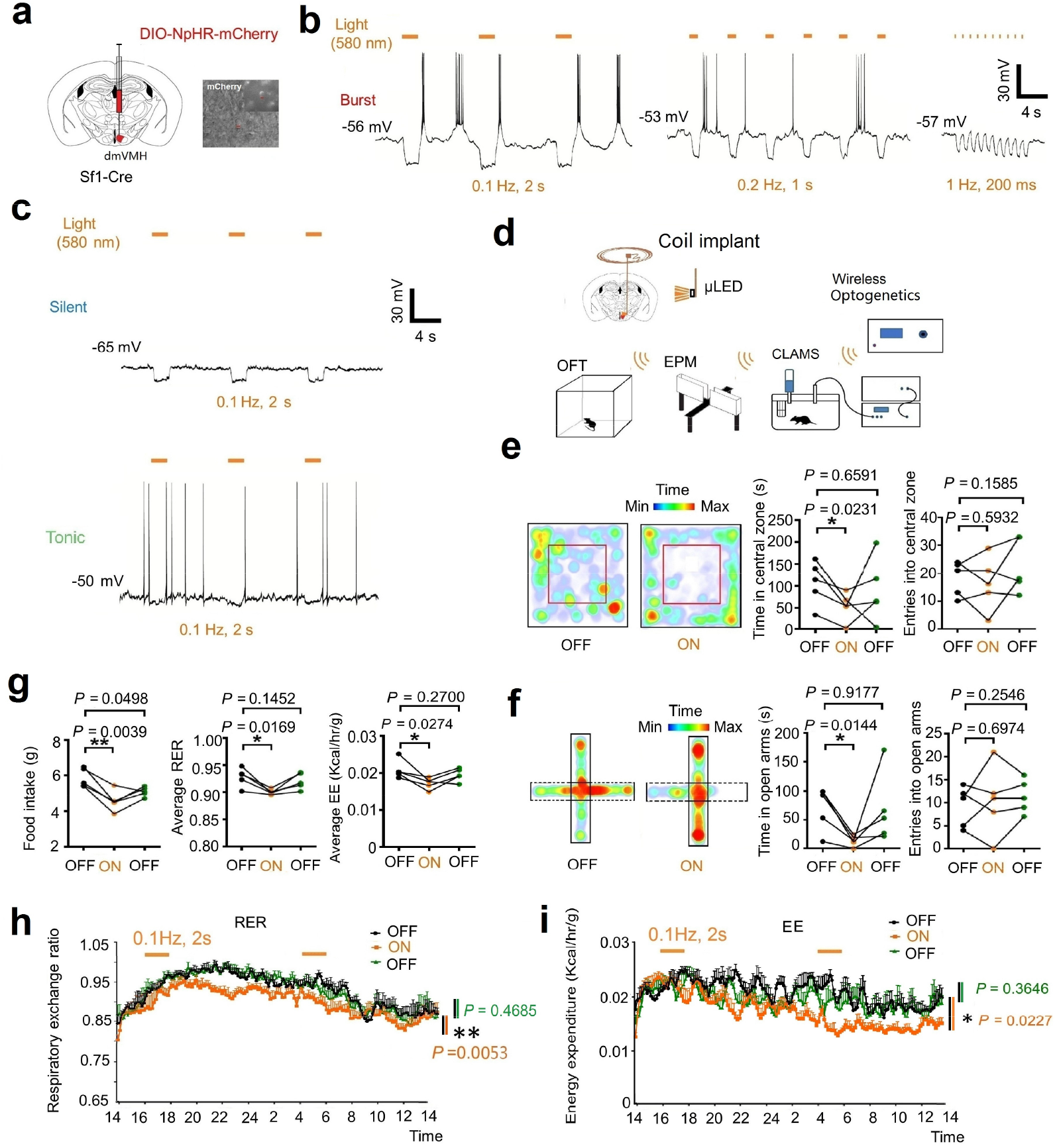
Optogenetic activation of burst firing neurons in dmVMH induced anxiety-like behavior and energy expenditure changes. **(a)** Schematic of dmVMH injection of NpHR AAV viral vector to induce burst firing *in vivo*. **(b)** Whole-cell recordings of yellow light-evoked burst firing in brain slices (yellow light: 590 nm, left: 0.1 Hz, 2 s; middle: 0.2 Hz, 1 s; right: 1 Hz, 200 ms), 0.1 Hz successfully induced activation of burst firing neurons. **(c)** 0.1 Hz and 2 s yellow light illumination exerted no significant influence on silent or tonic-firing dmVMH neurons. **(d)** Illustration of wireless optogenetic manipulation of dmVMH neurons and behavioral analysis in free-moving mice. **(e)** Open field test before, during, and after light illumination: residence time in central area decreased during 10-min yellow light illumination (n = 5, paired Student’s *t*-test, *P* = 0.0237), though no obvious changes were observed in number of entries into central area (paired Student’s *t*-test, *P* = 0.5932). **(f)** Elevated plus-maze test before, during, and after 5-min light illumination. Residence time in open arms decreased during yellow light illumination (n = 5, paired Student’s *t*-test, *P* = 0.0144) and recovered after light-off; no obvious changes were observed in number of entries into open arms (paired Student’s *t*-test, *P* = 0.6974). **(g)** Food intake and metabolism were monitored during optogenetic manipulation of dmVMH neurons in free-moving mice. Food intake decreased during light stimulation (paired Student’s *t*-test, *P* = 0.0039, n = 5 mice in each group), also average RER (paired Student’s *t*-test, *P* = 0.0169) and EE (paired Student’s *t*-test, *P* = 0.0274). **(h** and **i)** RER and EE curve shifted (two-way ANOVA, RER: *P* = 0.0053, *F*(1, 8) = 14.38; EE: *P* = 0.0227, *F*(1, 8) = 7.7918) when applying two yellow light stimulation trials (0.1 Hz, 2 s; for 2 h). Data are means ± SEM; * *P* < 0.05, ** *P* < 0.01.

We then applied low-frequency yellow light illumination to induce burst firing *in vivo* and performed behavioral and energy metabolic tests in NpHR-expressing mice (Fig. 3d). This led to decreased residence time in the central area of the open field and open arm of the elevated plus maze, mimicking the effects of chronic stress-induced anxiety-like behavior (Fig. 3e,f). We also applied wireless optogenetics to induce burst firing of neurons and simultaneously monitored the metabolism of mice using CLAMS. We applied 0.1-Hz yellow light illumination at the start of the test period for 2 h and repeated the stimulation after 12 h, with consecutive monitoring of mouse energy metabolism for 24 h. Results indicated that average RER, energy expenditure, and food intake in the test period decreased significantly compared with the baseline level before the test period (Fig. 3g-i). Taken together, our results indicated that enhancement of burst firing induced by local optogenetic manipulation in the dmVMH was sufficient to affect anxiety-like behavior and energy expenditure changes, which simulated the phenotypes of chronically stressed mice.

### Burst firing in dmVMH is mediated by T-VGCC

Bursting is an important firing pattern in neural systems and is essential for specific information transmission and function regulation^30^. The T-VGCC, including its three isoforms (Cav3.1, Cav3.2, and Cav3.3), is a pacemaker that can generate LTS ^26,31^. Unlike hyperpolarization-activated cyclic nucleotide-gated channel (HCN), the T-VGCC can induce higher frequency firing and has a threshold near to RMP. It has been reported that Cav 3.1 and Cav 3.2 alone display strong burst firing with a low-voltage threshold, whereas Cav3.3 contributes to burst firing in a different way ^31^. The expression of T-VGCC is highly correlated with T-type calcium currents, which directly affect the strength and width of bursting ^29,32^. Given the known role of T-VGCC in burst firing, we examined T-VGCC expression in the dmVMH after chronic stress.

Based on immunohistochemical analysis, Cav3.1, Cav3.2, and Cav3.3 were all found to be expressed in the dmVMH of naïve mice. The signals of Cav3.1 were much stronger than those of Cav3.2 and Cav3.3 (Fig. 4a). To investigate whether differential expression of the T-VGCC occurs after chronic stress, we performed acute dissection of dmVMH tissue from brain slices and harvested single dmVMH cells after patch recordings to extract RNA, then performed qRT-PCR to quantify T-VGCC expression. The expression of Cav3.1 in the dmVMH of stressed mice was significantly higher than that of the control group, whereas no obvious changes were observed in Cav 3.2 or Cav 3.3 at either the tissue or single-cell level (Fig. 4b-c).

**Fig. 4.**
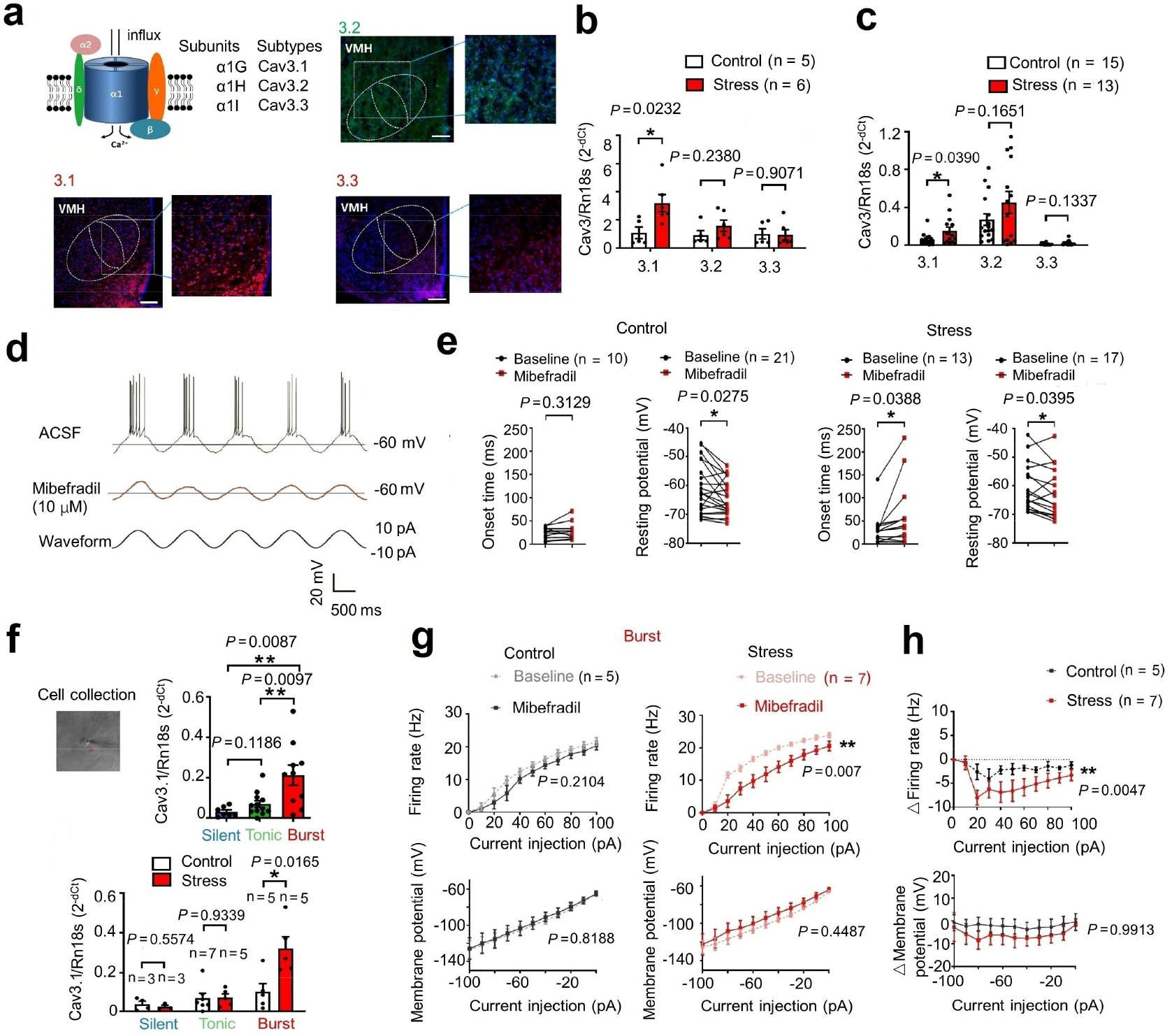
T-VGCC mediated enhancement of burst firing in dmVMH neurons under chronic stress. **(a)** Schematic of structure of T-VGCC located on cell membrane (left top). Representative immunofluorescence showing Cav 3.1 (left bottom), Cav 3.2 (right top), and Cav 3.3 (right bottom) expression in dmVMH, respectively, with stronger Cav3.1 expression observed. Scale bar is 200 μm. **(b)** Quantification of Cav 3.1, Cav 3.2, and Cav 3.3 expression in dmVMH tissue between control (n = 5 mice) and chronic stress groups (n = 6 mice). Expression of Cav 3.1 was significantly up-regulated under chronic stress conditions (unpaired Student’s *t*-test, *P* = 0.0232). **(c)** Single-cell qRT-PCR analysis of Cav 3.1, Cav 3.2, and Cav 3.3 expression in dmVMH neurons between control (n = 16 cells) and chronic stress groups (n = 14 cells). Expression of Cav 3.1 was significantly up-regulated under chronic stress conditions (unpaired Student’s *t*-test, *P* = 0.0390). **(d)** Evoked burst firing trace of dmVMH neurons without and with T-VGCC antagonist (mibefradil, 10 μM), 10 pA current injection was given in cosine waveform. **(e)** Effects of mibefradil on onset time and RMP of dmVMH neurons from wild-type (n = 15) and chronic stress groups (n = 13). Significant differences were observed in both onset time and RMP (paired Student’s *t-*test, *P* = 0.0388 and *P* = 0.0395) in the stress group, whereas the control group demonstrated obvious changes in RMP but not onset time (paired Student’s *t*-test, *P* = 0.0275 and *P* = 0.3125). **(f)** Single-cell qRT-PCR analysis of Cav 3.1 expression among three neuronal subtypes in dmVMH. Upper: burst firing neurons (n = 11) showed higher Cav 3.1 expression than other two subtypes (unpaired Student’s *t*-test, *P* = 0.0133 compared with silent neurons (n = 7), *P* = 0.0139 compared with tonic-firing neurons (n = 12)); bottom: Cav 3.1 expression in burst firing subtype showed significant differences between control and chronic-stress groups (unpaired Student’s *t*-test, silent: control, n = 3, stress, n = 3, *P* = 0.5574; tonic-firing: control, n = 7, stress, n = 5, *P* = 0.9339; bursting: control, n = 5, stress, n = 6, *P* = 0.0165). **(g)** Effects of mibefradil on suprathreshold and subthreshold activity in dmVMH burst firing neurons in control (n = 5) and chronic stress groups (n = 7). Mibefradil inhibited T-VGCC and caused a right frequency-current curve shift (two-way ANOVA, control, *P* = 0.2140, *F*(1, 8) = 1.854; stress, *P* = 0.0077, *F*(1, 12) = 10.22); lower, mibefradil application exerted no obvious influence on current-voltage curve of burst neurons (two-way ANOVA, control, *P* = 0.8188, *F*(1, 10) = 0.0573; stress, *P* = 0.4487, *F*(1, 10) = 0.6218). **(h)** Obvious differences were observed in suprathreshold activity (two-way ANOVA, *P* = 0.0047, *F*(1, 110) = 20.53), but not in subthreshold (two-way ANOVA, *P* = 0.9913, *F*(1, 33) = 4.958) membrane potential of dmVMH burst firing neurons between control (n = 5) and chronic stress groups (n = 7) after application of mibefradil. Data are means ± SEM. * *P* < 0.05, ** *P* < 0.01.

To further confirm the contribution of the T-VGCC in dmVMH burst generation, we applied mibefradil, an antagonist of T-VGCC, in whole-cell recording experiments. Results indicated that the burst firing of dmVMH neurons elicited by a 10-pA current injection was indeed inhibited (Fig. 4d). Furthermore, we found that application of mibefradil increased the onset time in the dmVMH neurons of the stress group, but not in the control group, but the change in RMP was similar between the two groups (Fig. 4e). Moreover, as we collected single neurons after whole-cell recordings for T-VGCC quantification, we combined the single-cell qRT-PCR and electrophysiological classification data to determine the differential expression of Cav3.1 in the dmVMH neuronal subtypes. Results showed that Cav3.1 expression was much more enriched in dmVMH bursting neurons, especially after chronic stress (Fig. 4f). Consistently, the Cav3.2 antagonist ascorbate ^33^ did not significantly affect burst firing of dmVMH neurons (Extended Data Fig. 3). We also investigated the effects of mibefradil on the frequency-current curves using the same stimulus protocol, and found that the firing rate of bursting neurons was decreased by bath application of mibefradil(*P* = 0.007, Fig. 4g, right), but the changes in the firing rate were much more significant in the stress group than in the control group (*P* = 0.0047, Fig. 4h, upper). These data support the idea that the T-VGCC, especially the Cav3.1 isoform, mediates burst firing in the dmVMH.

To investigate the *in vivo* effects of T-VGCC blockade, we bilaterally implanted a cannula and precisely delivered mibefradil or saline into the dmVMH of chronically stressed mice to block burst firing (Extended Data Fig. 4a). Results showed that mibefradil infusion increased the time spent in the central area of the open field and open arm of the elevated plus maze compared with the control group treated with saline (Extended Data Fig. 4b). We also investigated energy metabolic changes after oral administration of mibefradil, and found it to have no effect on food intake. However, administration of mibefradil significantly rescued the decreased RER found in stressed mice. Taken together, these data indicate that the T-VGCC in the dmVMH mediates chronic stress-induced behavioral and energy expenditure changes.

The function of the T-VGCC is tightly related to postsynaptic excitatory/inhibitory state ^34,35^. Several studies have suggested that the generation of burst firing mediated by T-VGCC activation in certain regions is highly dependent on membrane potential ^35,36^, and the NMDA receptor plays a critical role in regulating burst firing ^26^. Glutamate is a common excitatory neurotransmitter and its concentration around neurons may affect membrane potential ^37^. We applied different glutamate receptor antagonists into artificial cerebrospinal fluid (ACSF) during whole-cell recordings of burst firing neurons. Consistent with previous studies ^26^, our results indicated that blockade of NMDA receptors, but not AMPA receptors, significantly inhibited burst firing and caused membrane hyperpolarization, which made it more difficult to reach the T-VGCC threshold (Extended Data Fig. 5).

### Knockdown of Cav3.1 in dmVMH decreased burst firing and ameliorated chronic stress response

Given the important role of Cav 3.1 in mediating burst firing in the dmVMH, we next tested whether down-regulation of Cav 3.1 expression in the dmVMH was sufficient to ameliorate anxiety-like behavior and energy metabolic changes induced by chronic stress. We developed a lentivirus-mediated RNAi method to interfere with expression of Cav3.1 in the dmVMH under chronic stress, with the shRNA-expressing vector injected into the dmVMH bilaterally prior to chronic-stress exposure (Fig. 5a). To test the efficacy of shRNA-mediated Cav3.1 silencing, we used immunofluorescence to confirm the efficient knockdown of Cav3.1 in the dmVMH of stressed mice (Fig. 5b). The effects of Cav3.1 knockdown on burst firing were tested using whole-cell recording on brain slices obtained from Cav 3.1 knockdown mice. Results showed that 7 out of 17 dmVMH neurons from stressed mice displayed evoked burst firing, whereas only 2 out of 13 dmVMH neurons succeeded after Cav3.1 knockdown (Fig. 5b), suggesting burst firing inhibition induced by RNAi of Cav3.1.

**Fig. 5.**
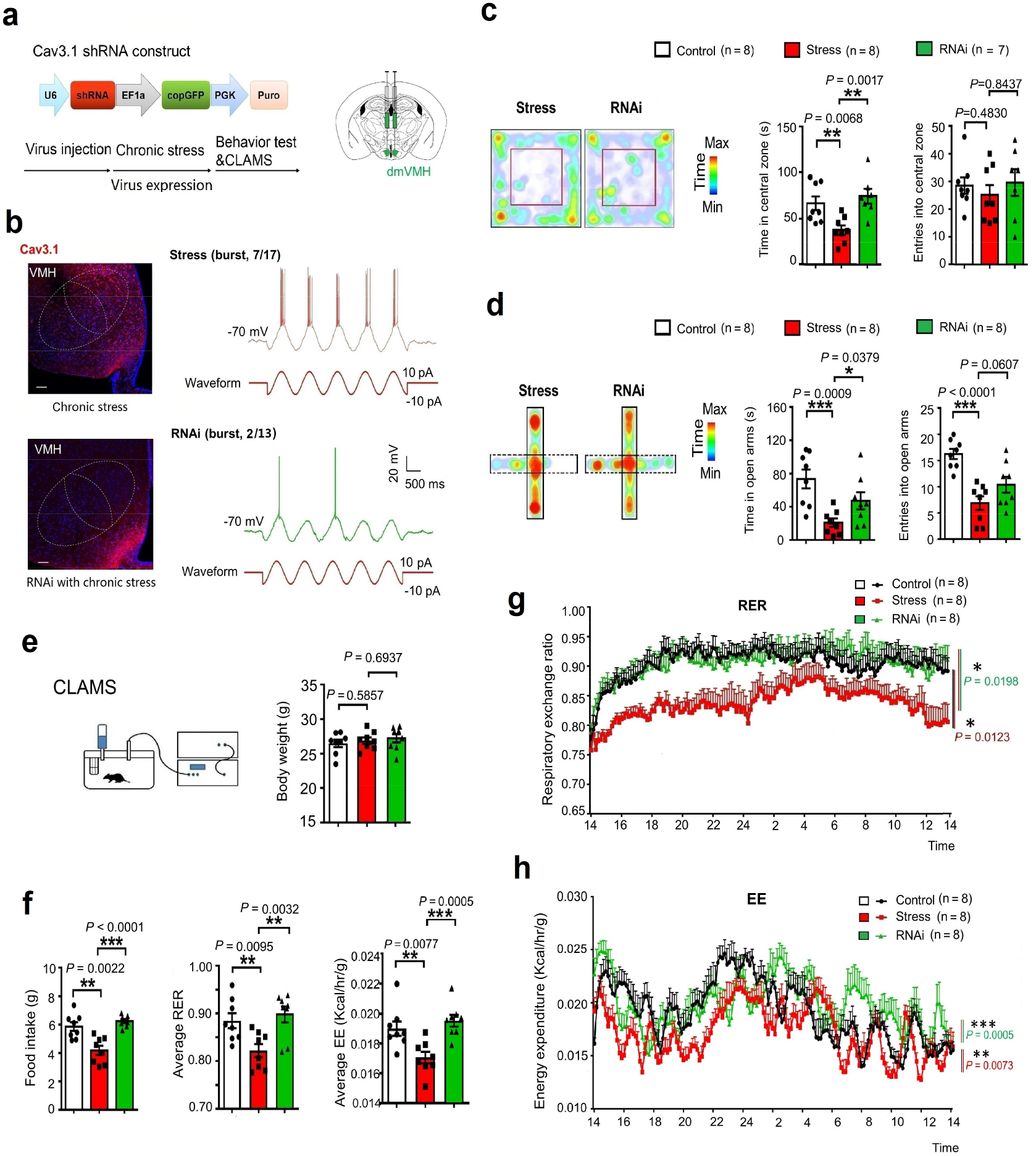
Knockdown of Cav3.1 in dmVMH decreased burst firing, and rescued anxiety-like behavior and metabolic alteration induced by chronic stress. **(a)** Schematic of Cav3.1 shRNA construct and injection of shRNA-expressing lenti-viral vector into dmVMH to interfere with Cav 3.1 expression. **(b)** Representative images of dmVMH Cav 3.1 immunostaining in chronic stress and RNAi (under chronic stress) animals (left). Proportion of burst firing neurons decreased in RNAi group (2/13, 15.38%) compared with stress group (7/17, 41.18%) (right). Scale bar is 100 μm. **(c)** Time spent in central area and number of entries into central area of open field in control, chronic stress, and RNAi groups. Residence time: control (n = 8) versus chronic stress group (n = 8), *P* = 0.0068; chronic stress versus RNAi group (n = 7), *P* = 0.0017. Number of entries: control versus chronic stress group, *P* = 0.4830; chronic stress versus RNAi group, *P* = 0.8437 (unpaired Student’s *t*-test). **(d)** Time spent in open arm and number of entries into open arm of elevated plus-maze in control, chronic stress, and RNAi groups. Residence time: control versus chronic stress group, *P* = 0.0009; chronic stress versus RNAi group, *P* = 0.0379. Number of entries: control versus chronic stress group, *P* < 0.0001; chronic stress versus RNAi group, *P* = 0.0607 (unpaired Student’s *t*-test, 8 mice in each group). **(e)** No significant differences in average body weights of control (n = 8), chronic stress (n = 8) and RNAi (n = 8) groups were observed after four weeks of chronic stress (unpaired Student’s *t*-test, control versus stress group, *P* = 0.5857; chronic stress versus RNAi group, *P* = 0.6937). **(f)** Food intake of mice in control, chronic stress, and RNAi groups: control versus chronic stress group, *P* = 0.0022; chronic stress versus RNAi group, *P* < 0.0001 (unpaired Student’s *t*-test, 8 mice in each group). Average RER of control, chronic stress, and RNAi groups (unpaired Student’s *t*-test: control versus chronic stress group, *P* = 0.0095; chronic stress versus RNAi group, *P* = 0.0032). Average EE of control, chronic stress, and RNAi groups (unpaired Student’s *t*-test: control versus chronic stress group, *P* = 0.0077; chronic stress versus RNAi group, *P* = 0.0005). **(g)** 24-h RER curve of control, chronic stress, and RNAi groups (two-way ANOVA: control versus chronic stress group, *P* = 0.0123, *F*(1, 14) = 8.244; chronic stress versus RNAi group, *P* = 0.0198, *F*(1, 13) = 7.055; 8 mice in each group). **(h)** 24-h EE curve of control, chronic stress, and RNAi groups (two-way ANOVA: control versus chronic stress group, *P* = 0.0005, *F*(1, 14) = 20.21; chronic stress versus RNAi group, *P* = 0.0073, *F*(1, 14) = 9.851; 8 mice in each group).Data are means ± SEM, * *P* < 0.05, ** *P* < 0.01, *** *P* < 0.001.

To investigate phenotypic changes in Cav3.1 knockdown mice, we carried out behavioral and energy metabolic tests after viral expression of shRNA of Cav3.1 for 4–5 weeks. Importantly, behavioral tests showed that Cav 3.1 knockdown significantly increased residence time and entries into the central area of the open field and open arm of the elevated plus-maze of stressed mice (Fig. 5c,d), suggesting that local knockdown of Cav 3.1 in the dmVMH was sufficient to rescue stress-induced anxiety-like behavior. For energy metabolic tests, we fasted all mice overnight and analyzed energy expenditure using CLAMS. Data showed that Cav 3.1 silencing effectively rescued the chronic stress-induced decrease in RER (Fig. 5f), as well as the reduction in food intake (Fig. 5f) and energy expenditure (Fig. 5a and h). No obvious body weight differences were observed among the control, stress, and RNAi groups (Fig. 5e).

Together with our optogenetic manipulation experiments, our results consistently demonstrated that Cav 3.1 in the dmVMH was both sufficient and necessary to elicit burst firing of dmVMH neurons. Furthermore, knockdown of Cav 3.1 in the dmVMH ameliorated chronic stress-induced phenotypic abnormalities, including anxiety-like behavior, lower RER, and decreased food intake.

## Discussion and conclusion

Previous studies have indicated that the dmVMH is an important stress coping center to balance aversive behavior and energy seeking ^2^ and may be a possible hub connecting stress-induced emotion and energy expenditure. Here, we reported that chronic stress induced burst firing enhancement in the dmVMH, which was critical for inducing anxiety-like behavior and energy expenditure alteration in stressed mice. Bursting is an important firing pattern in the central nervous system and is also important for specific information transmission. Based on our data, we determined that dmVMH burst firing neurons play an important role in connecting the emotional state of anxiety and energy homeostasis, and that T-VGCC Cav 3.1 is essential for dmVMH burst firing and contributes to chronic stress-induced behavior and energy expenditure changes (Figure S6).

### dmVMH is responsive in chronic stress

Chronic unpredictable stress is detrimental to both mental and physical health. Responses to chronic stress can vary from emotional disorders, like anxiety and depression, in the central nervous system to disturbed metabolism and energy homeostasis in peripheral organs ^3–5^. However, how these physiological processes integrate in the brain and the underlying neural mechanisms remain poorly understood. The dmVMH is implicated in integrating information in the limbic system and in maintaining energy balance ^38^. Furthermore, SF-1 neurons in the dmVMH constitute a nutritionally sensitive switch, which modulates the competing motivations of feeding and avoidance of environments full of acute stress ^2^. Given the important role of the dmVMH in balancing acute stress inputs and food-seeking, the effects of persistent unpredictable stress on the function of dmVMH neurons require further research.

In the current study, the dmVMH was found to be involved in integrating chronic stress inputs to regulate anxiety-like behavior and energy expenditure. We established an unpredictable and persistent stress mouse model and characterized the model with multiple energy metabolic, electrophysiological, and behavioral approaches. Consistent with previous reports ^3^, we found that unpredictable persistent stress induced aversive behavior, decreased food intake, and shifted the RER toward fat oxidation, without obvious effects on body weight. Furthermore, *in vivo* electrophysiology demonstrated that the power density of the theta band of local field potential was higher in the stress group than in the control group. To dissect the alteration in dmVMH neuronal activity under chronic stress conditions, we studied the electrophysiology of dmVMH neurons. Based on differences in electrical activity, we classified the dmVMH neurons into three subtypes: i.e., silent, tonic-firing, and bursting. By comparing these subtypes in the control and stress groups, we found an enhancement of burst firing in the dmVMH after chronic stress, including increased percentage of burst firing neurons, enhanced LFP, and decreased ISI of bursts. Using patch clamp recordings, our data demonstrated that chronic stressors indeed induced obvious electrophysiological changes in dmVMH neurons at the cellular level, further supporting the important role of the dmVMH in chronic stress.

### Burst firing of dmVMH is critical for chronic stress response

Action potentials that arrive in bursts provide more precise information than action potentials that arrive singly and further enhance the efficacy of neuromodulator release, thus revealing the special role of bursts in information transmission and processing ^30^. Many previous studies have indicated that burst firing of specific neuronal subpopulations is critical for performing specific functions. In the hippocampus, a single burst can produce long-term synaptic modifications ^39^. In the hypothalamus, burst firing is found in several regions, including the preoptic area and VMH ^40^. In the lateral habenula, bursting activity depends on the NMDA receptor and T-VGCC; and most importantly, can drive behavioral aversion and depression-like symptoms ^26^. To date, however, no specific study has reported on the functions of dmVMH burst firing under chronic stress. Here, we performed electrophysiological and gene expression studies and demonstrated that burst firing neurons in the dmVMH are actively involved in chronic stress response, and the ISI of burst firing was significantly correlated with the anxiety-like behavior in anxious mice after chronic stress.

To mimic burst firing *in vivo*, we applied optogenetic inhibition with an NpHR-expressing AAV vector, as optogenetic inhibition on the lateral habenula or hippocampal NpHR-expressing neurons can elicit burst firing in yellow light illumination intervals ^26,41^. However, burst firing in the dmVMH could not be elicited under an illumination frequency greater than 0.2 Hz in our electrophysiological studies, indicating regional specificity of T-VGCC activation kinetics. Importantly, we applied wireless optogenetics to successfully achieve burst manipulation in mice residing within a closed cage, followed by behavioral and energy metabolic analyses. Our data indicated that optogenetic-elicited burst firing was sufficient to promote the chronic stress-induced phenotypes described above. Taken together, these data show that VMH burst firing is an important hub for integrating and interacting anxiety-like emotional state and energy expenditure regulation.

### Mechanism of burst firing in dmVMH

The mechanism underlying neuronal burst firing is complicated and could explain specific functions in different brain regions. Previous studies have reported that various modulatory substances can promote depolarization of membranes and evoke burst firing of neurons. Oxytocin binds to its receptor in CA2 region and can activate the G protein-coupled pathway, and thus induce activation of several spike channels and closure of KCNQ potassium channels ^39^. Among the various mechanisms underlying burst initiation, activation of a low-threshold ion channel can induce burst firing in diverse brain regions ^26,31^. Typically, low-threshold T-VGCC is widely expressed in the central nervous system and can be activated by stimuli near the RMP to elicit burst firing. Molecular cloning has revealed three isoforms of T-type channel genes: i.e., Cav3.1, Cav3.2, and Cav3.3, which make distinct contributions to cellular electrical properties ^31,32,42^. Selective expression of Cav3.1 is sufficient to generate a strong rebound burst in deep cerebellar nuclear neurons, whereas expression of Cav3.2 or Cav3.3 alone does not generate a rebound discharge under normal conditions ^31^. Several studies have demonstrated that substitution of Cav3.1 or Cav3.2 for the native channel in model thalamic relay neurons causes elimination of high-frequency bursts, further confirming the role of Cav3.1 in burst firing ^32,42^.

Here, we identified the existence of Cav3.1, Cav3.2, and Cav3.3 in the dmVMH region. Cav3.1 and Cav3.2 displayed high expression in the dmVMH, whereas Cav3.3 showed lower expression, consistent with previous study ^40^. The T-VGCC antagonist mibefradil blocked burst firing of dmVMH neurons, thus suggesting the existence of T-VGCC-mediated burst firing in the dmVMH. Furthermore, combined analysis of electrophysiological classification and single-cell qRT-PCR, we found that Cav3.1 expression increased under chronic stress, which contributed to the enhancement of burst firing in the dmVMH. Furthermore, Cav3.1 demonstrated greater contribution to stress-induced enhancement of bursting in the dmVMH than the other isoforms.

We further investigated the necessity of Cav3.1 in chronic stress-induced anxiety-like behavior and energy metabolic disorders. We found that microinjection of mibefradil ameliorated anxiety-like behavior. Using RNAi methods, the expression of Cav3.1 in the dmVMH was specifically knocked down, resulting in the significant inhibition of neuronal bursting activity. Behavioral and energy expenditure experiments further demonstrated that the chronic stress responses described above were partially rescued. Thus, our findings consistently demonstrated that Cav3.1 may be a potential drug target for the treatment of anxiety and related energy metabolic disorders.

The current study has several limitations. Firstly, we focused on exploring neuronal activity changes in the dmVMH after chronic stress. As such, further research should focus on identifying the mechanism regulating Cav3.1 expression in the dmVMH, especially the receptors regulating Cav3.1 expression. Several factors might contribute to the increased percentage of burst firing neurons after stress, including stress-induced hormones or feeding-related peptides ^24^. Previous studies have reported that estrogen can regulate the expression and function of T-VGCC in vlVMH neurons, but not dmVMH neurons, through estrogen receptors ^40^. However, the mechanism underlying Cav3.1 expression influenced by hormones or neuropeptides after chronic stress needs further study.

Secondly, GABAergic neural circuits and astrocytes have been found in the VMH and are implicated in regulating energy expenditure ^43,44^, but their specific contributions to bursting activity need to be further investigated. We demonstrated that glutamate receptors (especially NMDA receptors) affect the generation of burst firing; however, whether synaptic glutamate uptake mediated by astrocytes affects burst firing requires further exploration. Moreover, as the function of the T-VGCC is highly membrane potential dependent, how upstream inputs integrate with the surrounding microenvironment to affect membrane potential, and hence regulate burst firing, needs to be further explored.

In summary, our study first identified bursting firing neurons in the dmVMH as a hub regulating emotional state and energy metabolic disorders. We also identified Cav 3.1 as the crucial regulator of bursting firing of dmVMH neurons. The results of this molecular and electrophysiological study should provide a more complete understanding of the chronic stress-induced emotional malfunction and peripheral metabolism disorders, and provides potential therapeutic targets for treating such malfunctions.

## Competing interests

The authors declare no competing interests.

## Acknowledgements

This project was partly supported by the National Natural Science Foundation of China (81471164, 31800881), Key Research Program of Frontier Sciences of Chinese Academy of Sciences (QYZDB-SSW-SMC056), and Shenzhen Governmental Basic Research Grant (JCYJ20170413164535041, JCYJ20180507182301299). We also thank Z.B Xu and B.F Liu for their help in transgenic mice husbandry and phenotyping. We are grateful to N.N Li and X.L Liu for the help in virus packaging.

## Author contributions

F.Y. and H.-L.H. conceived the idea and designed the experiments. J.S. D.-S.G and Y.-H.L performed all electrophysiological recordings, immunostaining, qRT-PCR and behavioral experiments, S.-P.C. and X.-Y.Z. helped with patch clamp recording and qRT-PCR, L.Z. helped with immunostaining and surgery, N.L. helped with the RNAi construction, Q.X. helped with in vivo electrophysiological recording, F.Y. and J.S. interpreted the results and wrote the manuscript with critical inputs from H.-L.H. and L.-P.W..

## Methods

### Animals

All procedures were carried out in accordance with protocols approved by the Ethics Committee of the Shenzhen Institutes of Advanced Technology, Chinese Academy of Sciences. Male C57BL/6 mice (4–8 weeks old) were purchased from the Guangdong Medical Laboratory Animal Center (Guangzhou, China). The SF-1-Cre mice (stock no: 012462) were obtained from Jackson Laboratories (Bar Harbor, ME, USA). Mice were housed at 22–25 °C on a circadian cycle of 12-hour light and 12-hour dark with ad-libitum access to food and water.

### Chronic stress procedures

All animals used in this study were male, and randomly assigned to either control or stress groups for chronic stress study. The stress group was daily subjected to one stressor which randomly chosen from following: (i) 10 mice squeezing in a relatively small cage (15 cm×10 cm×4 cm) for two hours. (ii) wet bedding in home cage overnight (200 ml water was added to moisten bedding). (iii) each mouse was tightly restraint in a tube for two hours. The chronic stress protocol lasted for 28 consecutive days. Mice in different stress groups received the same number of each stressor. Control animals were subjected to no stressors.

### Behavioral tests

#### Elevated plus maze (EPM)

Exposure (5-min) to EPM was used to assess locomotor activity and anxiety-related behaviors after chronic stress. The stress and control groups (n = 18–20/group) were placed in the center of the plus maze facing an open arm and behavior was recorded for the entire 5 min using an overhead-mounted camera. Videos recorded during the EPM test were analyzed with ANY-maze software (Stoelting Co., Wood Dale, USA) to acquire data on time spent in open and closed arms, locomotor activity (total distance travelled in maze), and entries into the open arm. Anxiety-related behavior is associated with less exploration in the open arm relative to overall exploration of all arms.

#### Open field test (OFT)

Open field exposure (10-min) was used to assess locomotor activity and anxiety-related behaviors after chronic stress. Mice (n = 18–20/group) were placed in the center of an open field and behavior was recorded for the entire 10 min. Videos were analyzed to acquire data on time spent in the center and corner areas, total locomotor activity, and number of entries into the center area. Anxiety-related behavior is associated with less exploration of the center area.

### CLAMS and energy expenditure

To characterize metabolic changes caused by chronic stress or Cav 3.1 knockdown, RER was measured by indirect calorimetry using a four-chamber open-circuit calorimeter (Oxymax Series; Columbus Instruments, Columbus, OH, USA). Mice were food deprived overnight, with body weight and chow in each cage weighed prior to the experiment. During the experiment, mice were housed individually in specially built Plexiglas cages (40 × 25 × 20 cm). Temperature was maintained at 22 °C with an airflow of 0.5 per min. Food and water were available *ad libitum*. Mice were subsequently monitored in the system for 24 h (whole light-dark cycle). Oxygen consumption (VO2) and carbon dioxide production (VCO2) were measured every 10 min. The RER was calculated as a quotient of VCO2/VO2, with 1 representing 100% carbohydrate oxidation and 0.7 representing 100% fat oxidation. Energy expenditure (kcal heat produced) was calculated as calorific value (CV) × VO2, where CV is 3.815 + 1.232 × RER ^3^. Metabolic data collected from the 24-h monitoring period were averaged for energy expenditure and RER. After the experiment, chow in each cage was weighed to calculate food intake.

### Slice preparation

Mice were deeply anesthetized with isoflurane and decapitated rapidly. Brains were then removed and transferred to chilled cutting solution within 3 min. Cutting solution contained (in mM): choline chloride 110; KCl 2.5; Na-pyruvate 0.6; MgCl_2_ 7.0; CaCl_2_ 0.5; NaH_2_PO_4_ 1.3; NaHCO_3_ 25; glucose 20 (pH 7.4). The chilled cutting solution was bubbled with carbogen (95% O2 and 5% CO2) for at least 30 min before use. Coronal slices (250–300 mm thick) were prepared on a vibratome (Series 1000, Warner Instruments, Berlin, Germany), and incubated in artificial cerebrospinal fluid (ACSF) containing (in mM): NaCl 125; KCl 2.5; Na-pyruvate 0.6; MgCl_2_ 1.3; CaCl_2_ 2.0; NaH_2_PO_4_ 1.3; NaHCO_3_ 25; glucose 10 (pH 7.4) bubbled with carbogen at 34 °C for 30 min. After incubation, all slices were equilibrated in ACSF at room temperature (24–26 °C) for at least 40 min. Single slices were placed on the recording chamber perfused with ACSF bubbled with carbogen at room temperature. Unless stated otherwise, drugs were applied with perfused ACSF.

### Electrophysiology

Whole-cell patch clamp recording was performed on VMH neurons. The dmVMH was identified based on landmarks (third ventricle). Recordings were obtained with multi-clamp 700B amplifiers (Molecular Devices, San Jose, USA) under visual guidance using a Nikon FN1 microscope (Tokyo, Japan). Electrophysiological data were acquired and analyzed using pClamp 10 software (Molecular Devices, San Jose, USA). Whole-cell recordings were performed with borosilicate glass electrodes (0.69 mm OD, 5–7 MΩ) with internal solution containing (in mM): K-gluconate 135.0; KCl 4.0; NaCl 2.0; HEPES 10; EGTA 4.0; Mg-ATP 4.0; Na-GTP 5.0. Osmolality was adjusted to 290–310 mOsm kg^-1^ with sucrose and pH was adjusted to 7.4 with KOH. After forming a high-resistance seal (GΩ), the cell was held in current-clamp mode for 7–10 min until access resistance stabilized. Resting membrane potential (RMP) was assessed at the beginning of the recording period after stabilization of access resistance, and periodically monitored throughout the recording by momentarily relieving the direct current injection. To elucidate differences among neurons, 800-ms current injections (−100 to 100 pA in 10 pA increments; 5 s interstimulus interval) were applied, and the number of action potentials evoked by each current injection, input resistance, and half-width of action potentials were determined.

### Drugs and reagents

All chemicals included in ACSF prescription were purchased from Sigma-Aldrich (Merck KGaA, Darmstadt, Germany). Mibefradil (40 μM) were purchased from Tocris (Bio-Techne, Minneapolis, USA) and used for electrophysiology and microinjection. NBQX (1,2,3,4-tetrahydro-6-nitro-2,3-dioxo-benzo[f]quinoxaline-7-sulfonamide, 30 μM) and AP5 (2-amino-5-phosphono-pentanoic acid, 30 μM) were acquired from Med Chem Express (MCE, Shanghai, China).

### Cell harvesting and single-cell Quantitative real-time polymerase chain reaction (qRT-PCR)

Single Cell-to-CT™ Kits (Thermo Fisher, USA) were used to perform single-cell qRT-PCR. After recording, single cells were collected and lysed to acquire total RNA. A Mastercycler 5333 PCR thermal cycler (Eppendorf, Germany) was used to perform reverse transcription and pre-amplification. TaqMan gene expression assay was used to assay each gene. Gene expression levels were normalized to the expression of housekeeping gene *Rn18s*. All protocols were performed according to the manufacturer’s instructions. qRT-PCR was performed with a Light cycler 480 (Roche, Switzerland).

### Tissue collection and qRT-PCR

To acquire dmVMH tissue, mice were first anesthetized with isoflurane. After bathing in cold 1% diethyl pyrocarbonate (DEPC) phosphate-buffered saline (PBS) solution, brains were acute cut on a vibratome. VMH tissues were dissected microscopically from those sections, and quickly transferred to TRIzol reagent. The manufacturer’s standard protocols for RNA extraction (TransGene, China) and synthesis (TOYOBO, Japan) were followed. Primers used for qRT-PCR included: Cav 3.1 (5’-TGG CCTTCTTCGTCCTGAAC-3’ and 5’-TTCTCCAGCCTCTTTAGTCGC-3’). Cav 3.2 (5’-CGGCCCTACT ACGCAGACTA-3’ and 5’-TTAAGGGCCTCGTCCAGAGA-3’), Cav 3.3 (5’-CTGCTATTCTCCAGCCCAGG-3’ and 5’-AGCTGCACCTCTTG CTTGT-3’). Expression of these gene was normalized to the expression of housekeeping gene β-Actin (*Act-b*).

### *In vivo* electrophysiology

Mice were implanted with two nickel-chromium wires (25-μm diameter; AM Systems, Sequim, USA) targeting the dmVMH, connector was bind to wires and fixed on skull with dental cement. After total recovery from surgery, mice were returned to the recording sessions. Neuronal activity was collected using a Plexon Multichannel Acquisition Processor system (Plexon, Dallas, USA). Local field potentials (digitized at 1 kHz sampling rate, low-pass filtered up to 250 Hz) were recorded simultaneously for 90 min with a gain of 5 000×. After the recording sessions, mice were anesthetized by pentobarbital sodium and perfused intracardially. The electrode recording position was marked by histological staining.

### Stereotaxic surgery and viral injection

For all stereotaxic surgery, 12–16-week-old mice were anesthetized by pentobarbital sodium (0.3% in saline, 1 ml/100 g, intraperitoneally) and placed in a stereotaxic apparatus for surgery. Stereotaxic surgical procedures were performed using standard protocols. To target the dmVMH, bilateral brain injection coordinates relative to bregma were chosen (AP, −1.58 mm; ML, ±0.3 mm; DV, −5.5 mm). Unless stated otherwise, 0.4 μL of viral vector was injected into the VMH at a rate of 0.1 μL/min using a 10-μL Hamilton syringe and a syringe infusion pump (World Precision Instruments, USA).

For RNAi knockdown study, Cav3.1-shRNA (U6-shRNA-EF1a-copGFP-PGK-Puro) was injected into the VMH with lenti-viral vector. Mice were housed for four weeks following injection for viral expression. Anxiety-like behavior and metabolic tests were performed at the end of the paradigm. To assess the knockdown effectiveness of Cav 3.1 shRNA, mice were perfused with 4% paraformaldehyde (PFA) and brain tissues were removed for immunostaining analysis after the final session.

For microinjection, a stainless-steel guide-cannula (0.6 mm outer diameter and 0.4 mm inner diameter, RWD, Shenzhen, China) was implanted into the diencephalon to target the VMH (AP, −1.58 mm; ML, ±0.3 mm; DV, −5.0 mm). The guide-cannula was fixed to the skull using dental cement and three stainless steel screws. Each guide-cannula was sealed with a stainless-steel wire for protection from obstruction. Before behavioral testing, the stainless-steel wire was replaced with an injection cannula (0.5 mm deeper than guide-cannula to break astrocyte aggregation), through which the drug dissolved in saline (400 nl) was delivered at 100 nl/min into the VMH. After microinjection, mice were rested for 30 min before tests.

### Wireless optogenetic manipulation

To achieve wireless optogenetic manipulation of burst firing neurons, AAV9-DIO-NpHR-mCherry was injected unilaterally into the VMH in the left hemisphere. Mice were housed for four weeks following injection for viral expression before initiation of experiments. A copper coil with a light emitting diode (LED) on the right side of the tip (Inper, Hangzhou, China) was implanted into the brain of NpHR-expressing mice, with the LED located 0.2 μm left of the dmVMH. The coil and indicator were fixed on the surface of the skull bone with Vetbond Tissue Adhesive (3M, USA). Charging the coil was achieved through antennas surrounding the cage. To initiate burst firing of dmVMH neurons, 550-nm yellow light stimulation was performed at 0.1 Hz (2 s/pulse) in the NpHR-mCherry group during the metabolic (one illumination trial/12 h, each trial lasted for 2 h) and behavioral tests (OFT, 10 min; EPM, 5 min).

### Immunohistochemistry

Mice were first anesthetized by chloral hydrate (10% in saline, 1 ml/100 g, intraperitoneally), then perfused with 0.01 M PBS and 50 mL of 4% PFA transcardially. Brains were dissected and post-fixed in 4% PFA overnight and then transferred to 30% w/v sucrose solution for cryoprotection until sinking. Sections from the entire anterior-posterior range of the VMH were stained using an antibody specific to Cav3.1 (1: 100, Alomone labs, Cat# ACC-021, RRID: AB_2039779), Cav3.2 (1: 100, Alomone labs, Cat#ACC-025, RRID: AB_2039781), or Cav 3.3(1: 100, Alomone labs, Cat# ACC-009, RRID: AB_2039783). Briefly, sections were washed, permeabilized in 0.1% Triton X-100/PBS for 15 min/three times, washed again, and blocked in 10% normal goat serum (NGS) (w/v)/0.1% Triton X-100/PBS for 30 min. Primary antibody was added, and sections were incubated overnight at 4 ℃. The following day, sections were washed with 0.01 mM PBS, incubated with incubated with secondary antibody (Alexa Fluor 594 Goat Anti-Rabbit, Cat#115-585-003, RRID: AB_2338059 or Alexa Fluor 488 Goat Anti-Rabbit, Cat#111-547-003 RRID: AB_2338058) in 1% NGS/0.1% Triton X-100/PBS for 1 h at room temperature, then washed, mounted, and cover slipped with mounting medium containing DAPI.

### Statistical analysis

Slice electrophysiological data were analyzed with pCLAMP (Molecular Device, San Jose, CA, USA). *In vivo* electrophysiology data were analyzed with NeuroExplorer 5.0 (Plexon, Dallas, USA). All data were imported into Prism 7 (GraphPad Software, La Jolla, CA, USA) and normality was assessed using D’Agostino-Pearson tests to verify the appropriateness of the following statistical analyses. Unless stated otherwise, the data are presented as means ± SEM. Statistical significance was determined using two-tailed unpaired Student’s *t*-tests when comparing two groups, paired Student’s *t*-test when comparing the effects of different treatments in the same group. When multiple measures were compared between groups (e.g., current-frequency curves), repeated measures two-way analysis of variance (ANOVA) with Bonferroni’s test was used. Differences were considered significant at *P* < 0.05.

To classify dmVMH neurons, unsupervised cluster analysis was performed with SPSS v19 (Chicago, IL, USA) using squared Euclidean distances. The parameters for cluster analysis were chosen based on their lack of linear correlation with each other. The following electrophysiological parameters were included for analysis: onset time, evoked firing rate, input resistance, and potential sag by H-current (Hyperpolarization-activated current). Resting potential were excluded from the parameters for cluster analysis because of their linear correlation with onset times. All electrophysiological parameters were converted into standardized z-scores before clustering.

## Supplemental Information

**Extended Data Fig. 1.**
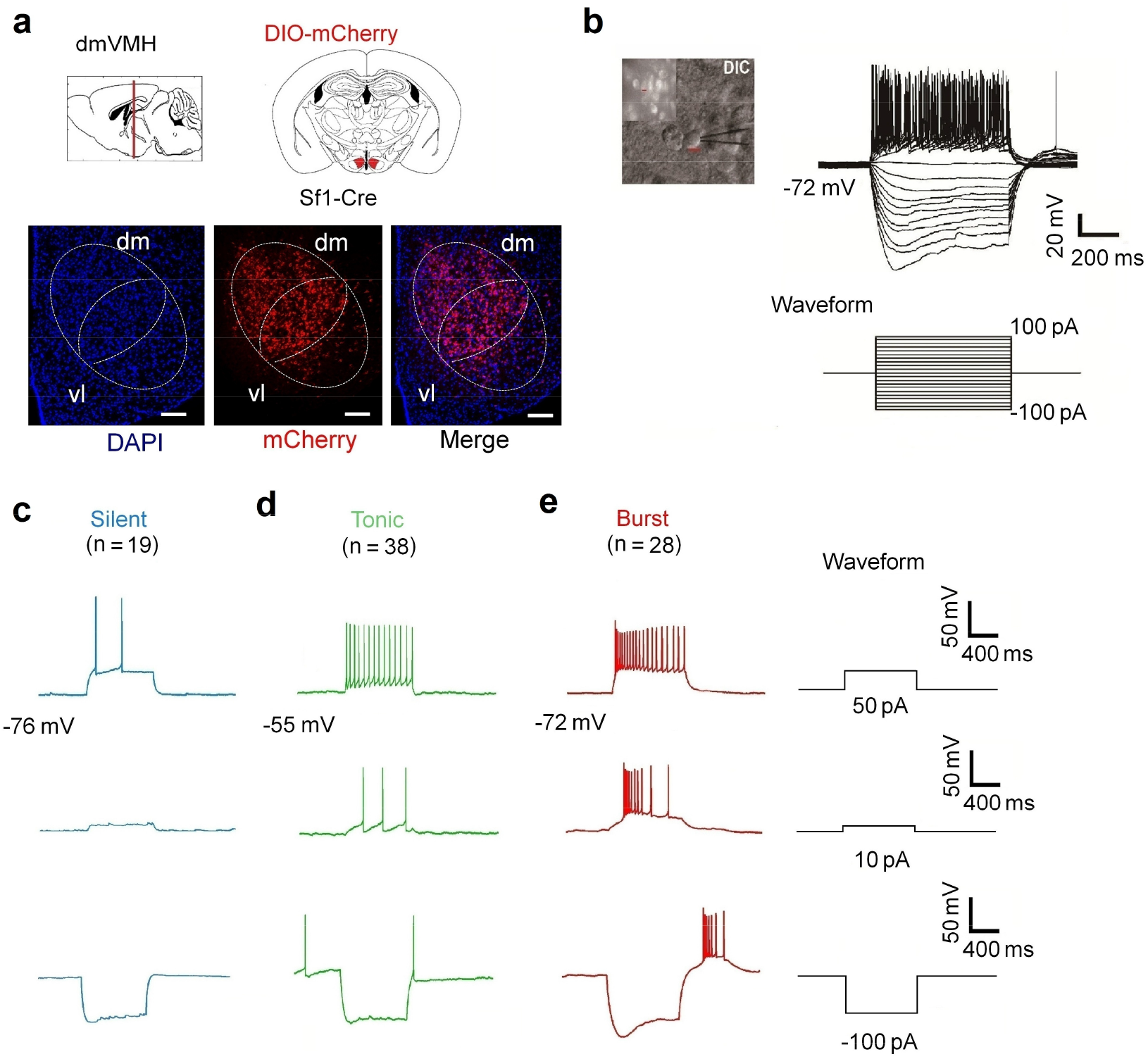
Electrophysiological properties of dmVMH neurons subtypes. Related to Fig. 2. (**a**) Schematic of location of dmVMH in coronal section slice of mouse brain, mCherry was specifically expressed in SF-1 (specific dmVMH marker) neurons. Scale bar is 300 μm. **(b)** Whole-cell recording trace from a dmVMH neuron, with current injection of −100 pA to 100 pA and 10 pA/step. (**c-e**), Representative traces of whole-cell recordings showing electrophysiological properties of silent (n = 19), tonic-firing (n = 38), and bursting (n = 28) dmVMH neuronal subtypes. Three subtypes exhibit different electrophysiological activity at 50, 10, and −100 pA current injection.

**Extended Data Fig. 2.**
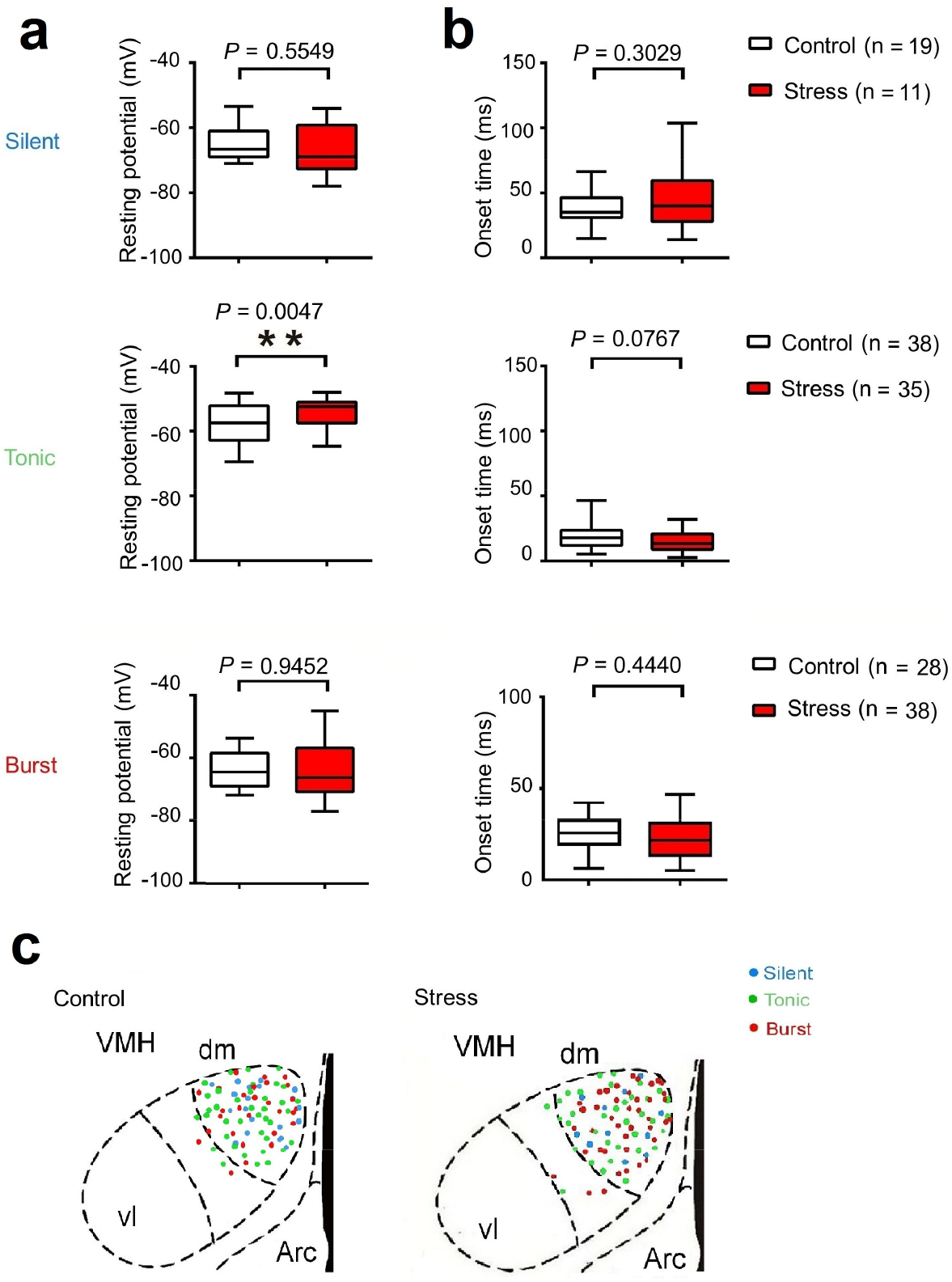
Electrophysiological comparison of three dmVMH neuronal subtypes in control and chronic stress groups. Related to Fig. 2. **(a)** Resting membrane potential (RMP) of three neuronal subtypes in control and stressed mice. Top: silent neurons in control (n = 19) and chronically stress mice (n = 13), *P* = 0.5549; Middle: tonic-firing neurons in control (n = 38) and chronically stress mice (n = 33), *P* = 0.0047; Bottom: bursting neurons in control (n = 29) and chronically stress mice (n = 38), *P* = 0.9452. **(b)** Onset time of three neuronal subtypes in control and stressed mice. Top: silent neurons in control (n = 19) and chronically stress mice (n = 13), *P* = 0.3029; Middle: tonic-firing neurons in control (n = 39) and chronically stress mice (n = 33), *P* = 0.0767; Bottom: bursting neurons in control (n = 27) and chronically stress mice (n = 38), *P* = 0.7704 (unpaired Student’s *t*-test). **(c)** Location of each recorded neuron in dmVMH of control and stressed group, no region specificity was observed among three subtypes. Data are means ± SEM; \**P* < 0.05, \*\**P* < 0.01. The box plotted at the median extending from the 25-75th percentile, and the whisker represents Min to Max distribution.

**Extended Data Fig. 3.**
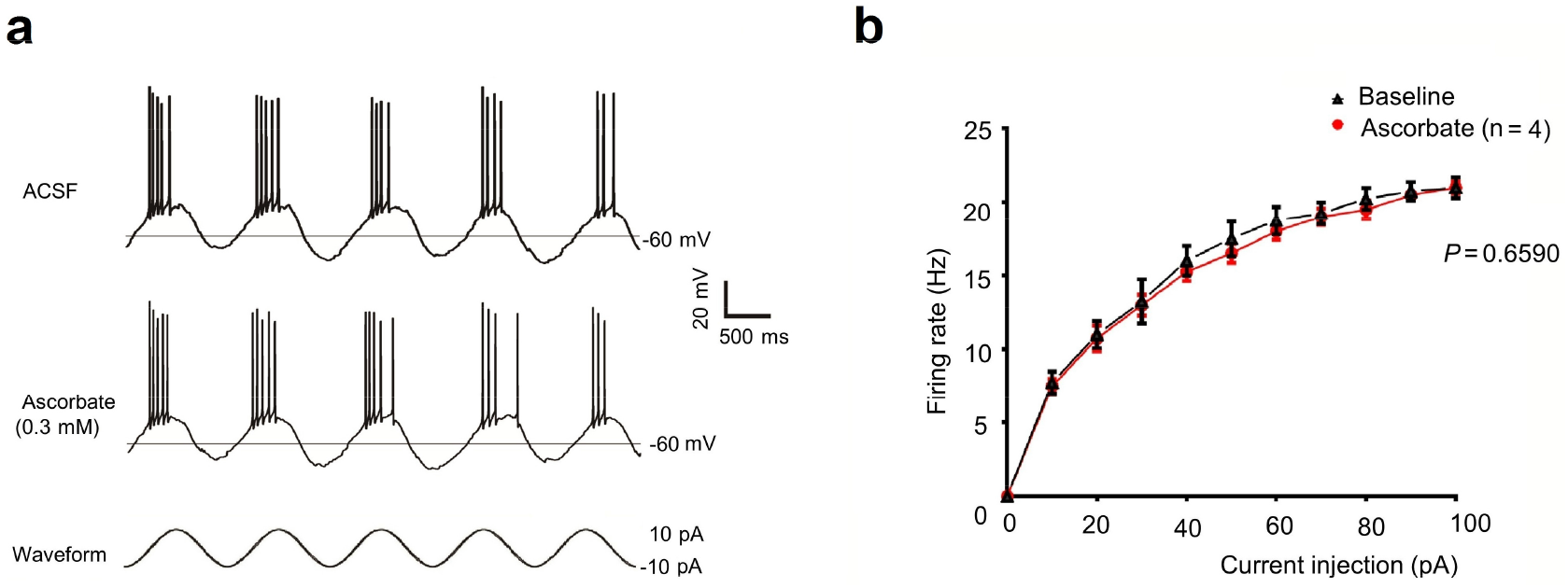
Ascorbate elicited no obvious effects on burst firing in dmVMH neurons. Related to Fig. 4. Ascorbate (antagonist of Cav3.2) did not block burst firing of dmVMH neurons (n = 4) induced by 10-pA current injection in cosine waveform. Ascorbate imposed no obvious influence on burst firing of dmVMH neurons, with no significant shifts found in frequency-current curve after ascorbate application (two-way ANOVA, *P* = 0.6590, *F*(1, 6) = 0.2152). \**P* < 0.05.

**Extended Data Fig. 4.**
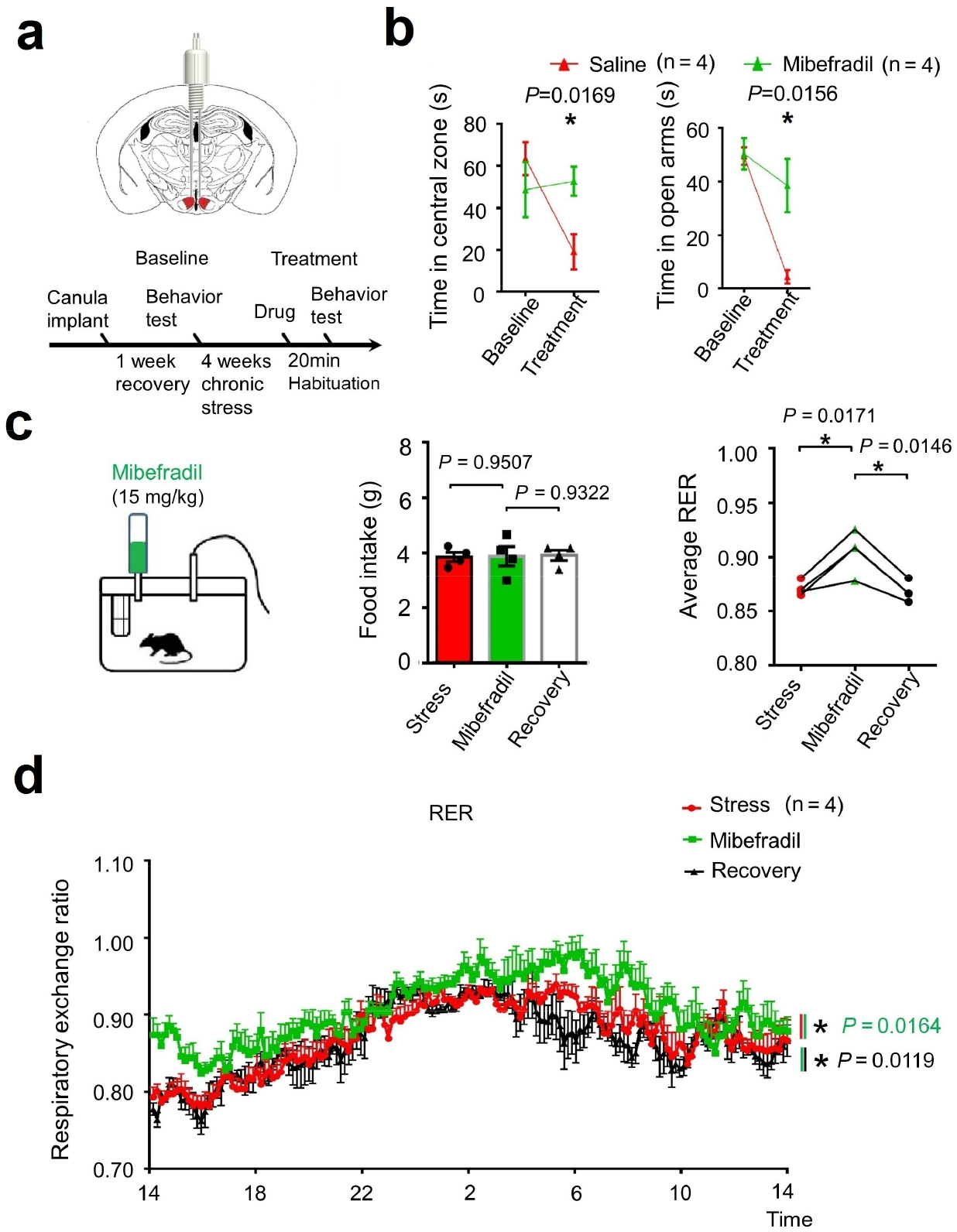
Injection of T-VGCC antagonist partially ameliorated anxiety-like behavior. Related to Fig. 4. **(a)** Schematic of microinjection of mibefradil in dmVMH through cannula. **(b)** Behavioral test before and after drug delivery; left, residence time in central area of open field of saline group decreased compared with mibefradil group (n = 4 in each group, *P* = 0.0169, unpaired Student’s *t*-test); right, residence time in open arm of saline group decreased compared with mibefradil group (n = 4 in each group, *P* = 0.0156, unpaired Student’s *t-*test). **(c-d)** Oral administration of mibefradil exerted no obvious change on food intake (n = 4 in each group, *P* = 0.9507, unpaired Student’s *t*-test) but increased average RER in stressed mice (*P* = 0.0171, unpaired Student’s *t*-test). Data are means ± SEM; \**P* < 0.05.

**Extended Data Fig. 5.**
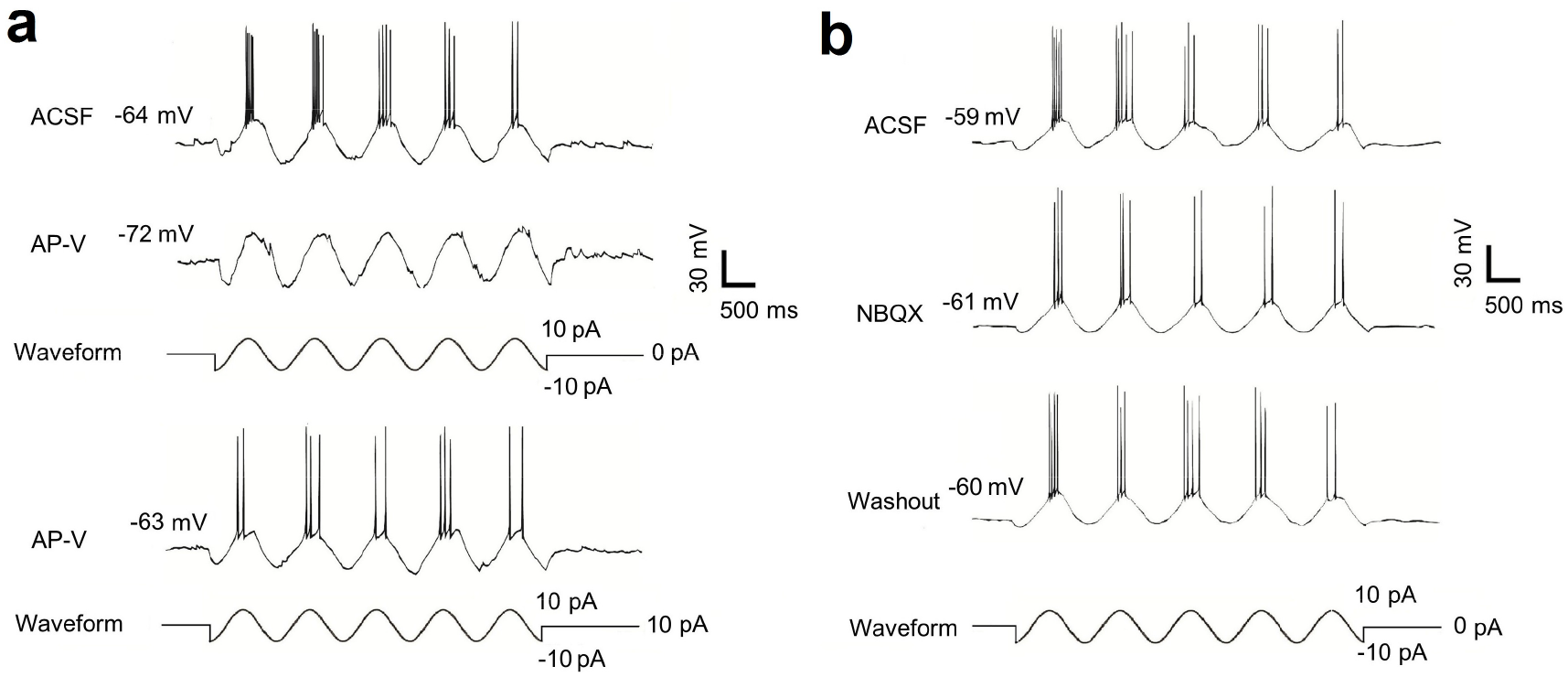
Glutamate receptor is critical for generation of burst firing in dmVMH. Related to Fig. 4. **(a)** Application of APV blocked generation of burst firing induced by cosine waveform current injection (−10 pA–10 pA, n = 4), even with depolarizing current injection (10pA) to eliminate the hyperpolarization caused by APV (lower). **(b)** Application of NBQX did not abolish generation of burst firing (n = 3).

**Extended Data Fig. 6.**
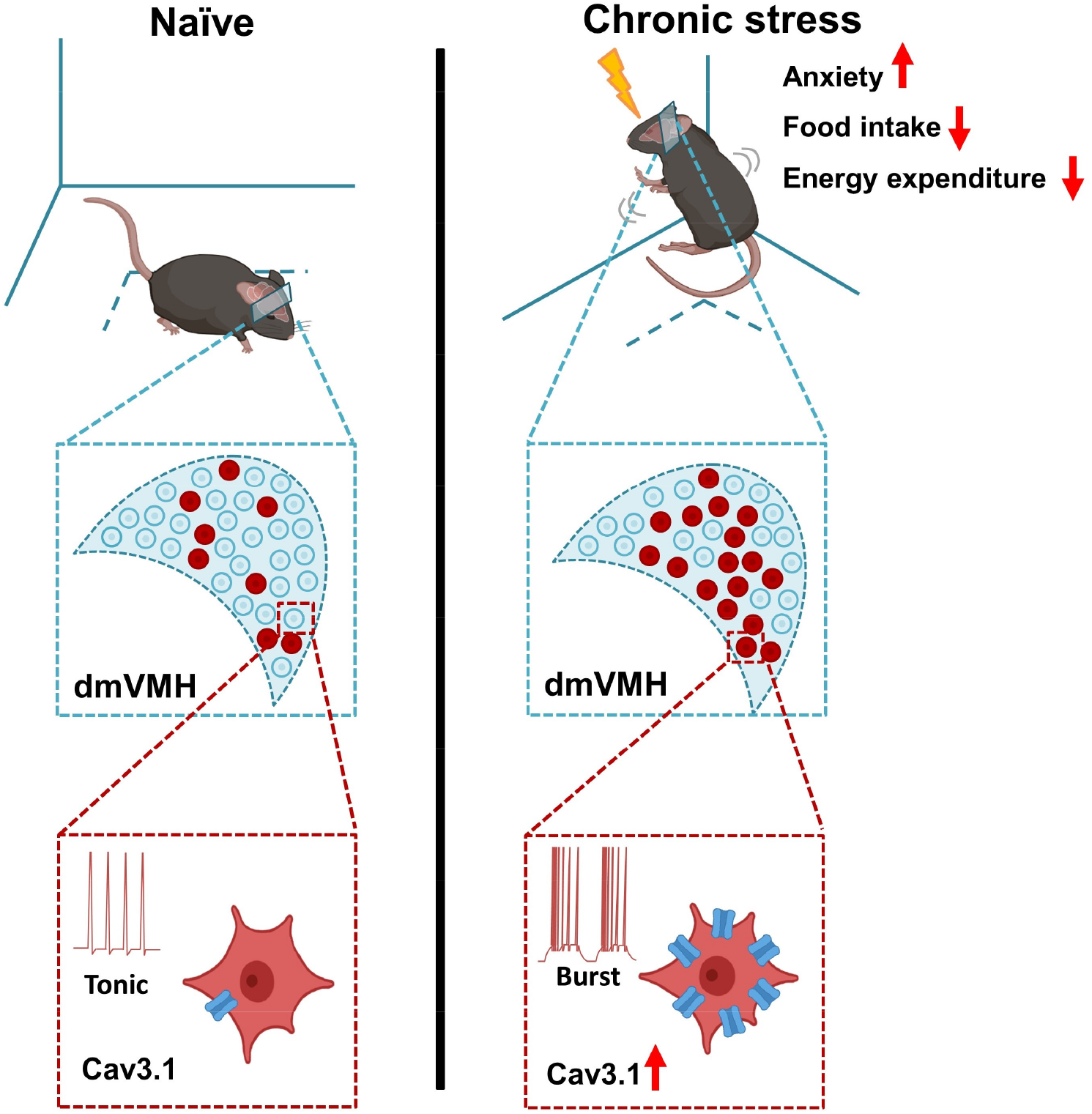
Schematic representation of the enhancement of burst firing in dmVMH neurons after chronic stress, which was mediated by the elevated expression of Cav3.1.

## Notes

### Competing Interest Statement

The authors have declared no competing interest.

